# Multiple clades of regulators contribute to bacterial phosphate homeostasis and pathogenesis

**DOI:** 10.1101/2025.09.08.674922

**Authors:** Caroline Vermilya, Eliot S. Joya Sandoval, Jana N. Radin, Gary J. Olsen, Bin Z. He, Thomas E. Kehl-Fie

**Affiliations:** Department of Microbiology and Immunology, University of Iowa, Iowa City, IA 52242 USA.; Department of Microbiology, University of Illinois at Urbana-Champaign, Urbana, IL 61801 USA; Carl R. Woese Institute for Genomic Biology, University of Illinois at Urbana-Champaign, Urbana IL 61801 USA; Department of Biology, University of Iowa, Iowa City, IA 52242 USA

**Keywords:** *Staphylococcus aureus*, PhoU, PhoPR, two-component system, phosphate homeostasis, regulation, virulence

## Abstract

Phosphate is both essential for life and toxic, necessitating the tight regulation of its acquisition. Based on *Escherichia coli*, most bacteria are thought to use a single accessory protein that monitors import to regulate phosphate homeostasis. This work reveals that most bacteria possess multiple distinct families of accessory regulators with each family regulating homeostasis in conjunction with a unique importer family. The antibiotic-resistant pathogen *Staphylococcus aureus* can obtain phosphate from divergent environments and possesses accessory-transporter pairs from all three identified groups. Investigations with *S. aureus* revealed that all three accessory proteins can regulate phosphate homeostasis, but that there is a hierarchy, which is dictated by the environment. Multiple accessory regulators are independently necessary for *S. aureus* to cause infection. Thus, microbes possess not one, but multiple distinct groups of accessory regulatory proteins and this diversity enables them to control phosphate homeostasis across environments, including those encountered during infection.

## Importance

Phosphate is both essential for life and toxic, necessitating the tight regulation of its acquisition. While previously thought to rely on a single accessory regulator, this work reveals that most bacteria possess mutiple distinct families of accessory regulators and that each family regulates homeostasis in conjunction with a unique importer family. Experiments utilizing *Staphylococcus aureus* revealed that all three accessory proteins can regulate phosphate homeostasis, but that an environment dependent hierarchy exists. Multiple accessory regulators are independently necessary for *S. aureus* to cause infection. Thus, microbes possess not one, but multiple distinct groups of accessory regulatory proteins and this diversity enables them to control phosphate homeostasis across environments, including those encountered during infection.

## Introduction

Phosphorous is required by all life, due to its critical role in nearly all biological processes. As inorganic phosphate (PO_4_^3-^ or P_i_), this irreplaceable nutrient is a component of nucleic acids, biological membranes, and has essential roles in signal transduction and energy metabolism (*1*). While needed, overaccumulation of P_i_ is toxic for both bacteria and eukaryotes (*2–17*). Thus, P_i_ uptake is tightly controlled (*1, 12, 18*). Despite the importance of P_i_ homeostasis and its intersection with numerous processes, our understanding of this critical mechanism in non-model systems and its contribution to the survival of microbes in their native environments is limited. As P_i_ uptake and homeostasis is linked with antibiotic resistance, bacterial virulence, and metal uptake, understanding how bacteria control their response to this nutrient will not only advance human health but also our ability to leverage microbes for environmental cleanup and biotechnological applications (*1, 19–24*).

The paradigm for P_i_ homeostasis in bacteria, *Escherichia coli*, uses a two-component signal transduction system (TCS), PhoBR, that does not directly sense P_i_. Instead, the activity of the sensor kinase PhoR is controlled by a single accessory protein, PhoU (*1, 4, 6, 25–29*). Classically, in conjunction with the PstSCAB P_i_-importer, PhoU is thought to sense the import of P_i_ and represses PhoBR activity when P_i_-flux is high (*1, 4, 6, 28, 29*). While many organisms possess homologs of PhoBR, PstSCAB, and PhoU, the *E. coli* system fails to capture the diversity of phosphate uptake and homeostasis machinery present across bacteria. Bacteria including *Vibrio fischeri* and *Staphylococcus aureus* possess a Na-P_i_ importer, NptA (*30, 31*). Furthermore, others including multiple Mycobacterial species, *Streptococcus pneumoniae* and *S. aureus* have multiple PhoU homologs (*9, 32–35*). How an expanded repertoire of accessory proteins impacts P_i_ homeostasis and the environments in which microbes can thrive is unknown.

*S. aureus* is an opportunistic pathogen that colonizes approximately 30% of the global population (*36*). Upon breaching the epithelium, it can infect virtually every host tissue, with the threat to human health growing due to widespread antibiotic resistance (*37, 38*). As with other pathogens, P_i_-importers and the TCS controlling P_i_ homeostasis, are critical to the ability of *S. aureus* to cause infection and influence antibiotic sensitivity (*22, 23, 27, 31, 35, 39*). In *S. aureus*, the PhoPR TCS controls P_i_ homeostasis, the expression of P_i_-importers, and is necessary for growth in P_i_-limited environments (*31, 35*). Differing from *E. coli*, *S. aureus* has three PhoU-like proteins or domain containing proteins, each encoded with a distinct P_i_-transporter: *pitR* with *pitA*, *phoU* with *pstSCAB*, and the C-terminal PhoU-domain of NptA (*35*). Given the importance of P_i_ homeostasis to bacterial physiology, pathogenesis, and antibiotic resistance, this work investigated the distribution of accessory regulatory proteins in bacteria and their contribution to P_i_ homeostasis and infection using *S. aureus*. These investigations revealed that PhoU-like proteins belong to three distinct clades, which each functioning in conjunction with a specific P_i_-transporter family to regulate P_i_ homeostasis. Further, these regulators were observed to work in a hierarchal manner with the hierarchy dictated by environment, resulting in multiple accessory proteins independently contributing to the ability of *S. aureus* to cause infection.

## Results

### Phylogenetic distribution of phosphate transporters and associated regulatory proteins

While the P_i_ homeostasis machinery present in *E. coli* does not capture the diversity present across microbes, a systematic analysis of these systems across bacteria has not been performed. To define the diversity of P_i_ uptake homeostasis machinery, 926 genomes from across the bacterial tree of life were evaluated for the presence of (*i*) the *S. aureus* P_i_-responsive TCS PhoPR (PhoP is PhoB in *E. coli*), (*ii*) homologs of the PstSCAB, PitA and NptA phosphate transporters, and (*iii*) PhoU-like proteins (PhoU, PitR and the PhoU-like domain of NptA). These proteins and their distributions in the genomes surveyed are detailed in Table S1 and Data S1. Homologs of PhoP and/or PhoR were identified in 698 (75.4%) of the genomes. Neither protein was identified in 228 genomes (comprised of diverse taxa including Aquificales, Bacteroidota, Planctomycetota, Chlamydiota, Rickettsiales, Campylobacterales, Spirochaetota, Mycoplasmatota). In 10 of the genomes, only one of these two proteins was identified. Identification of the PhoPR P_i_-responsive TCS in this analysis is the least certain, due to the existence of numerous paralogous two-component systems and selections are heavily biased toward genes that are genomically clustered with the genes for PhoU and/or the PstSCAB system. Thus, one possibility is that PhoPR homologs are present but sufficiently divergent from the quarry, preventing their identification. Alternatively, it is possible in the case of obligate intracellular genera, such as the Chlamydiota and Rickettsiales with reduced genomes, that this regulatory circuit has been lost.

Considering phosphate import, members of the Pst ABC-transporter family, represented by PstSCAB, were present most frequently, occurring in 767 (82.8%) of the genomes. The Pit phosphate inorganic transporter family, represented by PitA is present of 396 of the genomes (42.8%), while sodium-phosphate, NptA-type transporters were found in 306 (33.0%) of the genomes. A plurality of genomes 437 (47.2%) possess two P_i_-importer families, 327 (35.3%) possess only a single transporter family and 84 (9.1%) have members of all three transporter families. Notably 78 (8.4%) lacked an identifiable inorganic phosphate importer. This analysis underreports the diversity of transporters, as genomes were scored on the presence or absence of the transporter family and many possessed multiple copies of the same importer family (Data S1). Further, the NptA-type transporters are represented by 3 distinct annotations: NptA (exemplified by the *S. aureus* protein), NptA-PhoU (an NptA without the C-terminal PhoU-like domain, exemplified by the *B. subtilis* protein) and NptA.v (another NptA without a PhoU-like domain, exemplified by the *V. cholerae* protein), which were identified in 258 (27.9%), 38 (4.1%), and 39 (4.2%) of the genomes, respectively. Twenty-nine of the genomes have more than one of the NptA types. Also, NptA has numerous groups of proteins with unknown functions. Most notably “NptA-like protein YjbB” is present in about 150 of our genomes but is not sufficient for P_i_ import in *E. coli* (*40, 41*). Overall, these results indicate that there is much greater diversity in the mechanism utilized by bacteria to obtain P_i_ than previously appreciated or represented by the model systems studied to date_._

While the accessory proteins are broadly referred to as “PhoU”, an initial phylogenetic analysis suggests that PhoU and PitR were derived from an ancient gene duplication and have diverged since then, as shown by the PhoU-and PitR-related proteins forming distinct clusters in the gene tree and clustering graph (Fig. 1 and Fig. S1). This conclusion is supported by additional analyses with different subsets, demonstrating the gene tree is robust to sampling and tree reconstruction parameters (Fig. S1). One of the three accessory proteins was found in 823 (89%) of the genomes, with 445 (48%) possessing more than one. A PhoU homolog was identified in 722 (78.0%) of the genomes. Of the genomes with PhoU, 99.4% include a PstSCAB transporter, consistent with the function of PhoU being tied to PstSCAB. PitR is present in 359 (38.8%) of the genomes. When PitR is present, 97.8% of the time PitA is also present. With a sole exception, when both proteins are present, there is a *pitA* adjacent to *pitR* (some genomes have additional homologs that are not chromosomally clustered). In the genomes examined, 45 of them have a *pitA* adjacent to a gene that we annotated as “Uncharacterized membrane protein SCO1846” (named for its locus tag in the original *Streptomyces coelicolor* A3 (2) genome sequence). SCO1846 is 1/3 the size of PhoU and has no obvious similarity to it; it is noted because of its chromosomal location, and the observation that of the 45 PitA-containing genomes without PitR, 25 (54%) have a SCO1846 homolog. Of the 306 identified genomes with one of the three NptA-type transporters 258 (84%) possess at least one with a PhoU-domain. Cumulatively, this analysis suggests a tight linkage between each PhoU clade and a specific transporter family.

**Figure 1.**
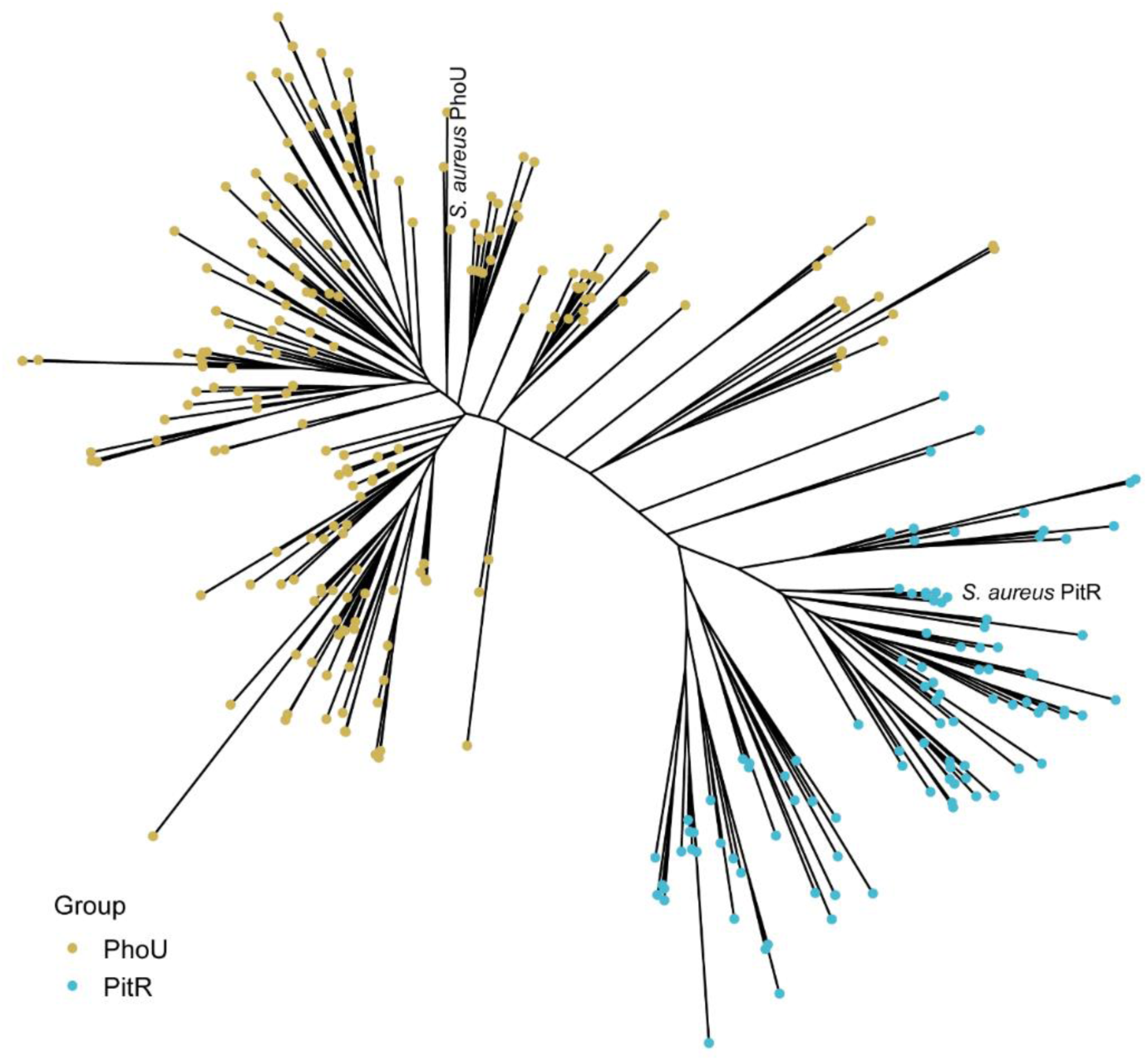
Unrooted phylogram shows the inferred evolutionary relationships between PhoU-and PitR-related proteins. PhoU-and PitR-related sequences were obtained from the InterPro database (Materials and Methods). After filtering by sequence identity at 70% and taxonomy at the genus level, 200 and 100 protein sequences were randomly chosen among the filtered representative sequences from each genus for both groups. Protein sequences were aligned using MAFFT v7 and a phylogenetic tree was reconstructed using RAxML v8.2.12 with LG+G substitution model. Branch lengths in the unrooted tree are proportional to the sequence divergence. *S. aureus* PhoU and PitR sequences are labeled.

### The PitR and NptA clades of accessory regulators control activation of phosphate-responsive two-component signal transduction systems

To date only members of the PhoU group of accessory regulators are known to modulate P_i_ homeostasis. However, loss of the staphylococcal PhoU does not alter activation of PhoPR (*35*). This combined with the current bioinformatic analysis led to the hypothesis that PitR or the PhoU-domain of NptA control phosphate homeostasis. As a first step in testing this hypothesis, Δ*pitR* and *nptA*_Δ*phoU*_ mutants were generated. The *nptA*_Δ*phoU*_ mutant, which lacks the C-terminal PhoU-domain, was created by deleting the nucleotides that encode the PhoU-domain from the chromosomal copy of *nptA*, thereby preserving native regulation. Deletion of the PhoU-domain does not eliminate the ability of NptA_ΔPhoU_ to import P_i_, as Δ*pstSCABnptA*_Δ*phoU*_ retains the ability to grow in alkaline (pH 8.4) PFM9 (phosphate-free M9-based medium) with a limiting concentration (50 µM) of P_i_ (Fig. S2) (*31*). Surprisingly, when excess P_i_ is present in an alkaline medium, the growth of Δ*pstSCABnptA*_Δ*phoU*_ is significantly less than the wild-type (Fig. S2), suggesting that the PhoU-domain of NptA may regulate import activity.

Next the activity of PhoPR was evaluated in wild-type, Δ*pitR,* Δ*phoU*, and *nptA*_Δ*phoU*_ in neutral (pH 7.4) P_i_-excess (5 mM), -intermediate (500 µM), and -limited (50 µM) medium, by assessing *pst* and *nptA* expression. In P_i_-limited medium, *pst* and *nptA* are highly, and with some minor variation, similarly expressed across all strains (Fig. 2). In P_i_-intermediate medium, *pst* expression is induced to similar levels in wild-type, Δ*pitR,* Δ*phoU* and *nptA*_Δ*phoU,*_ but to an extent less than in P_i_-limited medium (Fig. 2). The expression of *nptA* is also elevated in all strains following growth in P_i_-intermediate medium (Fig. 2). However, loss of PitR or the PhoU-domain of NptA results in increased *nptA* expression when compared to wild-type bacteria (Fig. 2). This is reversed by expression of either PitR or wild-type NptA from a plasmid (Fig. 2). In P_i_-excess medium, expression of *pst* and *nptA* in Δ*pitR,* but not Δ*phoU* or *nptA*_Δ*phoU*_, is elevated relative to wild-type (Fig. 2). The increased expression of the *pst* loci is reversed by ectopic expression of *pitR* or *nptA* (Fig. 2). These data indicate that the non-PhoU accessory regulators contribute to PhoPR activation and that PitR is the dominant regulator in *S. aureus*.

**Figure 2.**
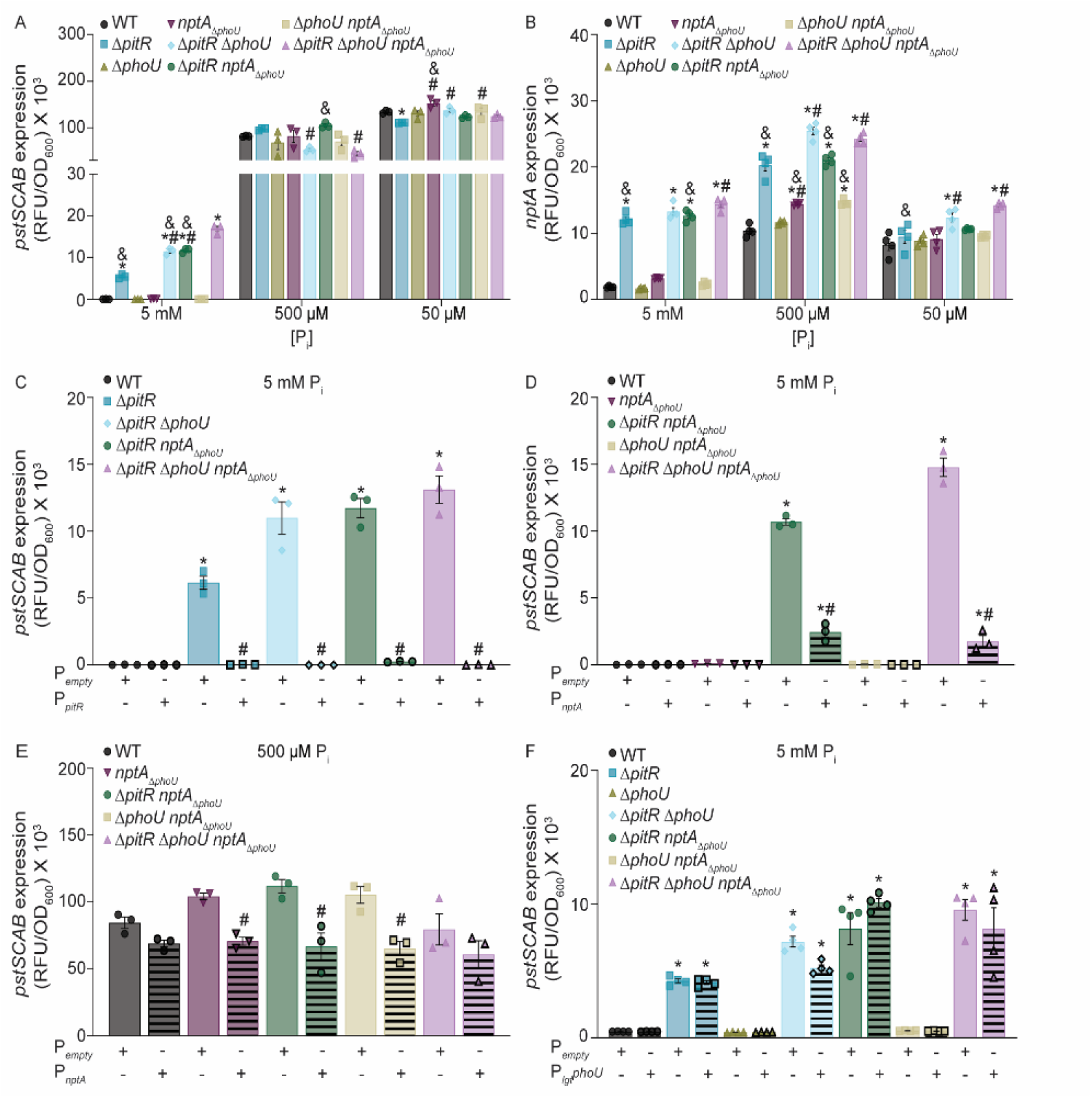
The PitR and NptA clades of accessory regulators control activation of phosphate-responsive two-component signal transduction systems. (A & B) *S. aureus* wild-type and the indicated strains containing (A) P*_pstS_-yfp* or (B) P*_nptA_-yfp* reporters were grown in PFM9, pH 7.4, supplemented with 5 mM, 500 µM, or 50 µM P_i_. The expression of *pstSCAB* or *nptA* was assessed by measuring fluorescence at T = 8 h. *\** = *P* ≤ 0.05 relative to wild-type at the same P_i_ concentration via one-way ANOVA with Tukey’s posttest. *#* = *P* ≤ 0.05 relative to Δ*pitR* at the same P_i_ concentration via one-way ANOVA with Tukey’s posttest. *&* = *P* ≤ 0.05 relative to Δ*pitR*Δ*phoUnptA*_Δ*phoU*_ via one-way ANOVA with Tukey’s posttest. *n* ≥ 3. Error bars = SEM. (C, D, E, F) *S. aureus* wild-type and the indicated strains containing the P*_pstS_-yfp* reporter and an empty vector (P*_empty_*), (C) P*_pitR_*, (D, E) P*_nptA_*, (F) P*_lgt_phoU* plasmids were grown in PFM9, pH 7.4, supplemented with 5 mM (C, D, F) or 500 µM (E) P_i_. The expression of *pstSCAB* or *nptA* was assessed by measuring fluorescence at T = 8 h. “+” designates the presence of the indicated plasmid, while “-” designates its absence. *\** = *P* ≤ 0.05 relative to wild-type at the same P_i_ concentration via one-way ANOVA with Tukey’s posttest. *#* = *P* ≤ 0.05 relative to the parent strain carrying P*_empty_* at the same P_i_ concentration via one-way ANOVA with Tukey’s posttest. *n* ≥ 3. Error bars = SEM.

### The three clades of regulators combine to regulate phosphate homeostasis

In P_i_-excess medium, *pst* expression in Δ*pitR*, is less than that of wild-type in P_i_-limited medium (Fig. 2), suggesting that additional regulators contribute to phosphate homeostasis. To evaluate the hypothesis that PhoU and the PhoU-domain of NptA are these regulators, a panel of double and triple PhoU homolog mutants (Δ*pitR*Δ*phoU*, Δ*pitRnptA*_Δ*phoU*_, Δ*phoUnptA*_Δ*phoU*_, and Δ*pitR*Δ*phoUnptA*_Δ*phoU*_) was created. Surprisingly, the loss of all three PhoU homologs in multiple independent clones does not result in an overt growth defect in TSB, a phosphate-replete medium (Fig. S3). This differs from *E. coli*, in which PhoU mutations are lethal and lead to the accumulation of compensatory mutations in PhoBR and PstSCAB (*4, 6, 28*). Considering this possibility, the *phoPR*, *pitA*, *nptA*, and *pstSCAB* loci were sequenced in Δ*pitR*Δ*phoUnptA*_Δ*phoU*_ and no mutations were observed. In *S. aureus*, phosphate-related growth defects can become apparent following growth in a phosphate-defined medium (*31*). Therefore, the growth of the mutant panel was evaluated in PFM9 supplemented with limiting, intermediate, or excess P_i_ at neutral pH. In the P_i_-intermediate and -limiting medium, all the strains grow similarly (Fig. 3 & S4). However, in the P_i_-excess medium, Δ*pitR*Δ*phoU* and Δ*pitR*Δ*phoUnptA*_Δ*phoU*_ display an extended lag phase but reach a final optical density similar to that of wild-type (Fig. 3). The growth defect of Δ*pitR*Δ*phoU* and Δ*pitR*Δ*phoUnptA*_Δ*phoU*_ in P_i_-excess medium is fully complemented by ectopic expression of *pitR* and partially complemented by ectopic expression of *phoU* (Fig. 3 & S4), indicating that the growth defect is due to loss of PitR and PhoU. These results indicate that in contrast to *E. coli* and other organisms, loss of PhoU-like regulation is not lethal to *S. aureus* (*4, 6–10, 28, 42, 43*).

**Figure 3.**
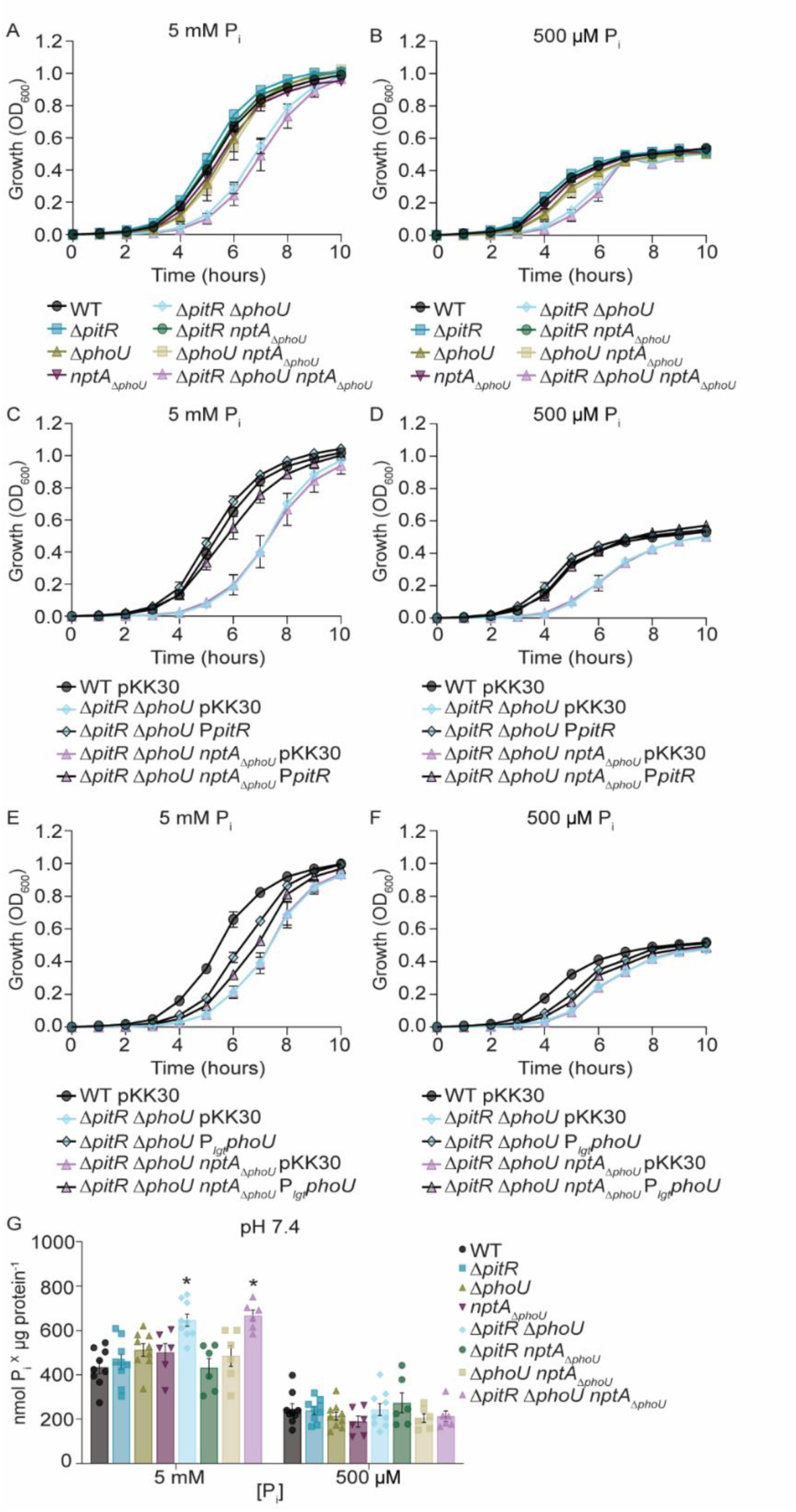
The staphylococcal PhoU homologs are not essential for viability. (A-F) *S. aureus* wild-type and the indicated strains were grown in PFM9, pH 7.4, supplemented with 5 mM or 500 µM P_i_. Growth was assessed by measuring absorbance at OD_600_. *n* ≥ 3. Error bars = SEM. C-F) Wild-type *S. aureus* and the indicated mutants containing an empty vector (pKK30), P*_pitR_* (C-D) or P*_lgt_phoU* (E-F) expression constructs were grown in PFM9 supplement with 5 mM or 500 µM P_i_ as indicated. n ≥ 3. Error bars = SEM. (G) *S. aureus* wild-type and the indicated strains were grown in PFM9, supplemented with 5 mM or 500 µM P_i_, at pH 7.4 and intracellular P_i_ was assessed. *\** = *P* ≤ 0.05 relative to wild-type at the same P_i_ concentration via two-way ANOVA with Dunnett’s posttest. *n* ≥ 5. Error bars = SEM.

P_i_ toxicity is due to excessive accumulation (*2–17*). The viability of Δ*pitR*Δ*phoUnptA*_Δ*phoU*_ in phosphate-replete media raises the possibility that loss of the staphylococcal accessory proteins does not impact cellular P_i_ levels despite the elevated transporter expression in Δ*pitR.* Therefore, P_i_ levels in the single, double, and triple PhoU mutants were assessed following growth in P_i_-intermediate and P_i_-excess medium at a neutral pH. In P_i_-intermediate medium, all the strains accumulate similar levels of P_i_ regardless of the pH (Fig. 3). However, in P_i_-excess medium both Δ*pitR*Δ*phoU* and Δ*pitR*Δ*phoUnptA*_Δ*phoU*_ accumulate more P_i_ than wild-type (Fig. 3). This indicates that the accessory regulators combine to regulate P_i_ homeostasis and suggests that microbes have inherently different abilities to cope with dysregulation of phosphate homeostasis and elevated cytoplasmic P_i_ levels.

### Multiple accessory proteins regulate phosphate homeostasis in a hierarchical manner

The submaximal expression of *nptA* and *pst* in Δ*pitR* and elevated cellular P_i_ in Δ*pitR*Δ*phoU* and Δ*pitR*Δ*phoUnptA*_Δ*phoU*_ lead to the hypothesis that PhoU and the PhoU-domain of NptA could also modulate the activity of PhoPR. To test this hypothesis, the expression of *pstS* and *nptA* was evaluated in the panel of double and triple mutants in PFM9 supplemented with limiting, intermediate, or excess P_i_ at neutral pH and compared to wild-type bacteria and Δ*pitR*. In both P_i_-limited and -intermediate medium, apart from small differences, *pst* expression is similar between all strains (Fig. 2) In contrast and unexpectedly, in P_i_-limited medium Δ*pitR*Δ*phoU* and Δ*pitR*Δ*phoUnptA*_Δ*phoU*_ have increased *nptA* expression when compared to wild-type or Δ*pitR.* (Fig. 2). At an intermediate-P_i_ concentration, all the double and triple mutants have elevated *nptA* expression when compared to wild-type, with expression in Δ*pitR*Δ*phoU* and Δ*pitR*Δ*phoUnptA*_Δ*phoU*_ exceeding that of Δ*pitR*. These observations suggest that reduced phosphate availability is not sufficient to maximally induce NptA expression.

In P_i_-excess medium, any strain lacking *pitR* has elevated expression of both *pst* and *nptA* when compared to wild-type (Fig. 2). However, Δ*pitRnptA*_Δ*phoU*_, Δ*pitR*Δ*phoU* and Δ*pitR*Δ*phoUnptA*_Δ*phoU*_ also express *pst* at a higher level than Δ*pitR,* with the triple mutant also having higher expression than the double mutants. A similar pattern occurs with *nptA*, except that only the triple mutant has elevated expression when compared to Δ*pitR.* Ectopic expression of *pitR, nptA,* or *phoU* reverses the overexpression of *pst* (Fig. 2). While PitR fully complements the double and triple mutants, the expression of NptA and PhoU return *pst* expression to a level similar to that of Δ*pitR*. Together these observations indicate that all three accessory protein families can modulate PhoPR activity but that they control P_i_ homeostasis in a hierarchical manner.

### Accessory regulators control homeostasis in conjunction with their cognate transporter

The clustering of each accessory regulator family with a specific transporter leads to the hypothesis that each is only capable of regulating phosphate homeostasis in conjunction with their cognate transporter. To test this hypothesis, *pstS* expression was assessed in wild-type, Δ*pitR*, Δ*phoU*, Δ*pitR*Δ*phoU*, Δ*pitRnptA*_Δ*phoU*_, Δ*phoUnptA*_Δ*phoU*_, and Δ*pitR*Δ*phoUnptA*_Δ*phoU*_ possessing a constitutively expressed copy of PhoU. Following growth in P_i_-replete media, expression of *phoU* does not compensate for the loss of PitR in any of the tested strains (Fig. 2). However, it does reduce *pst* expression in Δ*pitR*Δ*phoU* to a level comparable to Δ*pitR,* indicating that PhoU is functional. These results indicate that accessory regulators work with their cognate transporter to regulate phosphate uptake.

### Environment dictates the contribution of accessory proteins to controlling P_i_ homeostasis

The three P_i_-transporters possessed by *S. aureus* expand the environments in which it can thrive, with PitA promoting growth in acidic environments and NptA in alkaline environments (*31*). In the host, pathogens are faced with acidity within phagolysosomes, while blood and skin may alkalinize during bacterial infection, with pH influencing both the expression and functionality of the three *S. aureus* P_i_ transporters (*31, 44–46*). This leads to the hypothesis that the contribution of the accessory regulators may differ depending on the environment. To test this hypothesis, growth of the single, double, and triple mutants as well as *pst* and *nptA* expression was assessed in acidic (pH 6.4) and alkaline (pH 8.4) medium. Similar to a neutral medium, an *S. aureus* strain lacking all three accessory regulators is viable in both acidic and alkaline medium across a range of P_i_ concentrations (Fig. S5 & S6). However, growth of Δ*pitR*Δ*phoU* and Δ*pitR*Δ*phoUnptA*_Δ*phoU*_ was notably worse in alkaline media. As in neutral medium, the simultaneous loss of PitR and PhoU does result in an extended lag phase that can be complemented (Fig. S5 & S6). The Δ*pitR*Δ*phoUnptA*_Δ*phoU*_ mutant also accumulates more P_i_ than wild-type in acidic P_i_-replete medium (Fig. 4). P_i_ accumulation of Δ*pitR*Δ*phoU* and Δ*pitR*Δ*phoUnptA*_Δ*phoU*_ could not be assessed in alkaline medium due to growth defects. Taken together, these data indicate that staphylococcal phosphate acquisition is influenced by all three PhoU proteins in an environment-dependent manner.

**Figure 4.**
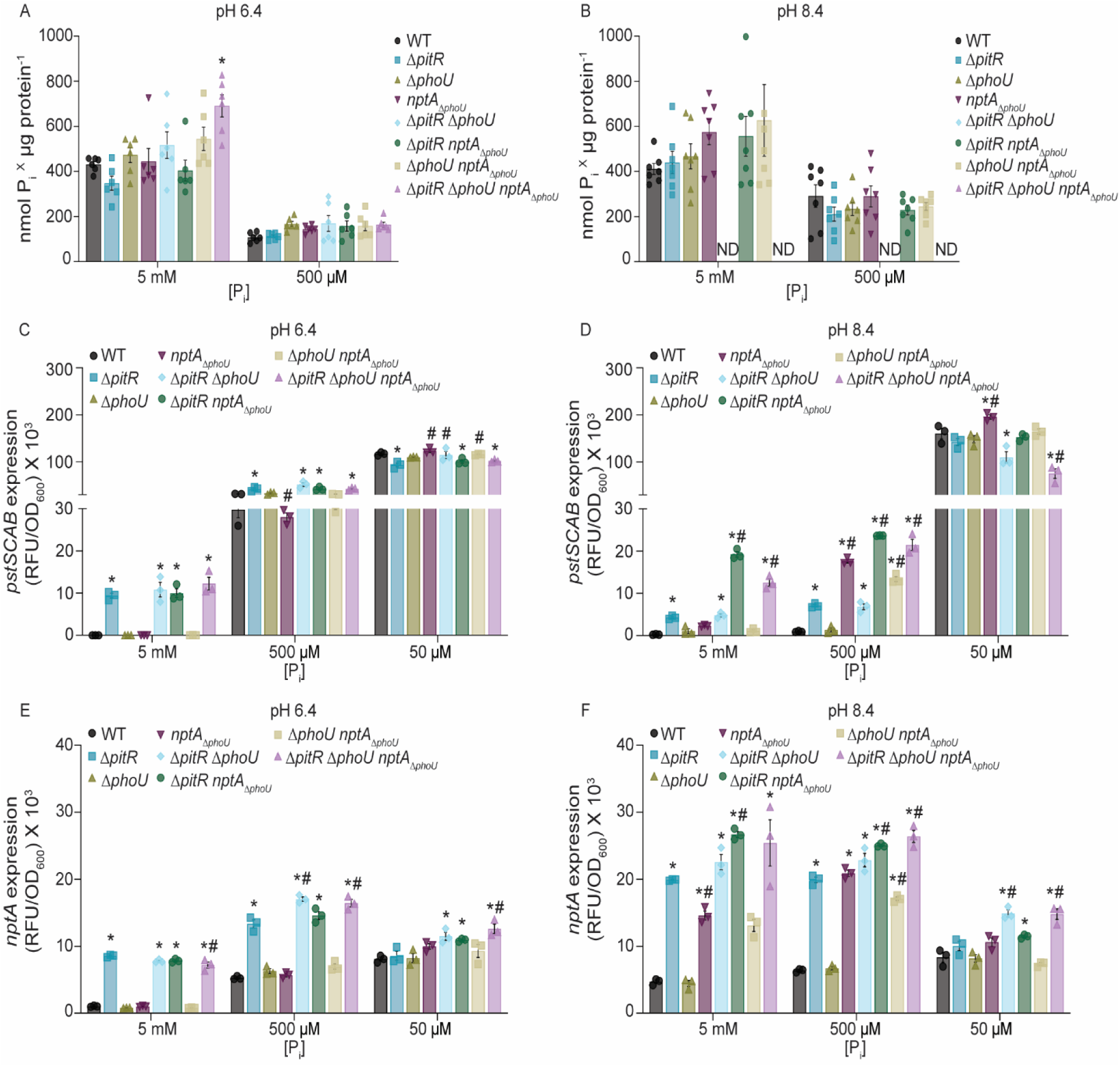
Environment dictates the contribution of accessory proteins to controlling P_i_ homeostasis. (A, B) *S. aureus* wild-type and the indicated strains were grown in PFM9, supplemented with 5 mM or 500 µM P_i_, at pH 6.4 (A) or 8.4 (B), and intracellular P_i_ was assessed. *\** = *P* ≤ 0.05 relative to wild-type at the same P_i_ concentration via two-way ANOVA with Dunnett’s posttest. *n* ≥ 5. Error bars = SEM. ND, not determined due to an inability to generate sufficient biomass. (C, D, E, F) *S. aureus* wild-type and the indicated strains containing (C & D) P*_pstS_-yfp* or (E & F) P*_nptA_-yfp* reporters were grown in PFM9, at pH 6.4 (C & E) or 8.4 (D & F), supplemented with 5 mM, 500 µM, or 50 µM P_i_. The expression of *pstSCAB* or *nptA* was assessed by measuring fluorescence at T = 8 h. *\** = *P* ≤ 0.05 relative to wild-type at the same P_i_ concentration via one-way ANOVA with Tukey’s posttest. *n* ≥ 3. Error bars = SEM. *#* = *P* ≤ 0.05 relative to Δ*pitR* at the same P_i_ concentration via one-way ANOVA with Tukey’s posttest.

To test the hypothesis that environment dictates the accessory regulator hierarchy, expression of *pst* and *nptA* was assessed in acidic and alkaline medium with limited, intermediate, and excess concentrations of P_i_. In P_i_-limited or -intermediate acidic medium, the pattern of *pst* expression in wild-type, the single, double, and triple mutants is similar to that observed in neutral medium (Fig. 2 & 4). Differing from neutral media with excess P_i_ at an acidic pH, the double and triple mutants do not express *pst* at a higher level than Δ*pitR*. In alkaline medium, the pattern of *pst* expression differs substantially from that observed in neutral medium. Most notably, in the P_i_-excess medium, loss of either PitR or the PhoU-domain of NptA greatly increases *pst* expression. Additionally, in medium with an intermediate P_i_ concentration, loss of the NptA PhoU-domain results in greater *pst* expression than in either wild-type bacteria or Δ*pitR*. In low P_i_ medium, *pst* expression is largely similar across all the strains. The altered expression of *pstSCAB* in acidic and alkaline medium is reversed upon ectopic expression of *pitR*, while ectopic expression of *nptA* reversed phenotypes observed in alkaline medium (Fig. S7).

Similar to *pst*, in acidic medium across all three P_i_ concentrations, the overall pattern of *nptA* expression in wild-type and the mutants is similar to that observed in a neutral medium (Fig. 2 & 4). However, *nptA* expression is muted overall. In contrast, in alkaline medium the overall expression of *nptA* increases, suggesting that its expression is influenced by pH. As with *pst* expression, loss of the PhoU-domain of NptA in P_i_-excess medium is sufficient to induce elevated *nptA* expression. The loss of this domain also results in a more profound impact on *nptA* expression in P_i_-intermediate medium. Cumulatively, these results demonstrate that the environment impacts P_i_ homeostasis and the respective contributions of the accessory regulators to controlling P_i_ homeostasis.

### Multiple accessory regulators contribute to the pathogenesis of *Staphylococcus aureus*

PhoPR and P_i_ importers are necessary for *S. aureus* to cause systemic infection, indicating that the host can present as a P_i_-limited environment (*31, 35*). However, the basal P_i_ level in human plasma ranges from 0.8 to 1.4 mM (*47*), suggesting that *S. aureus* is also exposed to excess P_i_ during infection. Considering this, the contribution of the three staphylococcal PhoUs to infection was evaluated. For these investigations, a systemic murine model of infection was used (*48*). Initially, C57BL/6 mice were infected with *S. aureus* wild-type, Δ*pitR*, Δ*phoU* and *nptA_ΔphoU_*. Similar bacterial burdens are recovered from the liver and heart of mice infected with wild-type and *nptA*_Δ*phoU*_ (Fig. 5). In contrast, 100-1000-fold fewer bacteria are recovered from these tissues following infection with Δ*pitR* or Δ*phoU* (Fig. 5). The capacity of the double and triple accessory protein mutants to cause infection was also assessed. Consistent with the loss of *pitR* or *phoU*, all double and triple mutants lacking either PitR or PhoU have decreased virulence (Fig. 5). In the liver and heart, when compared to wild-type, 1,000-10,000-fold fewer bacteria are recovered following infection with Δ*pitR*Δ*phoU*, Δ*pitRnptA*_Δ*phoU*_, Δ*phoUnptA*_Δ*phoU*_, and Δ*pitR*Δ*phoUnptA*_Δ*phoU*_ (Fig. 5). The double and triple mutants also display greater attenuation than the single mutants. In the liver, fewer bacteria are recovered following infection with Δ*phoUnptA*_Δ*phoU*_ than Δ*phoU*. Similarly, fewer bacteria are recovered following infection with Δ*pitR*Δ*phoUnptA*_Δ*phoU*_ than both Δ*pitR* and Δ*phoU* (Fig. 5). In the heart, fewer bacteria are recovered following infection with Δ*pitR*Δ*phoU* than Δ*phoU* (Fig. 5). Altogether, these data reveal that the accessory regulators both independently and collectively contribute to controlling phosphate homeostasis during infection and that hyperactivation of the P_i_ starvation response is detrimental to the pathogenesis of *S. aureus*.

**Figure 5.**
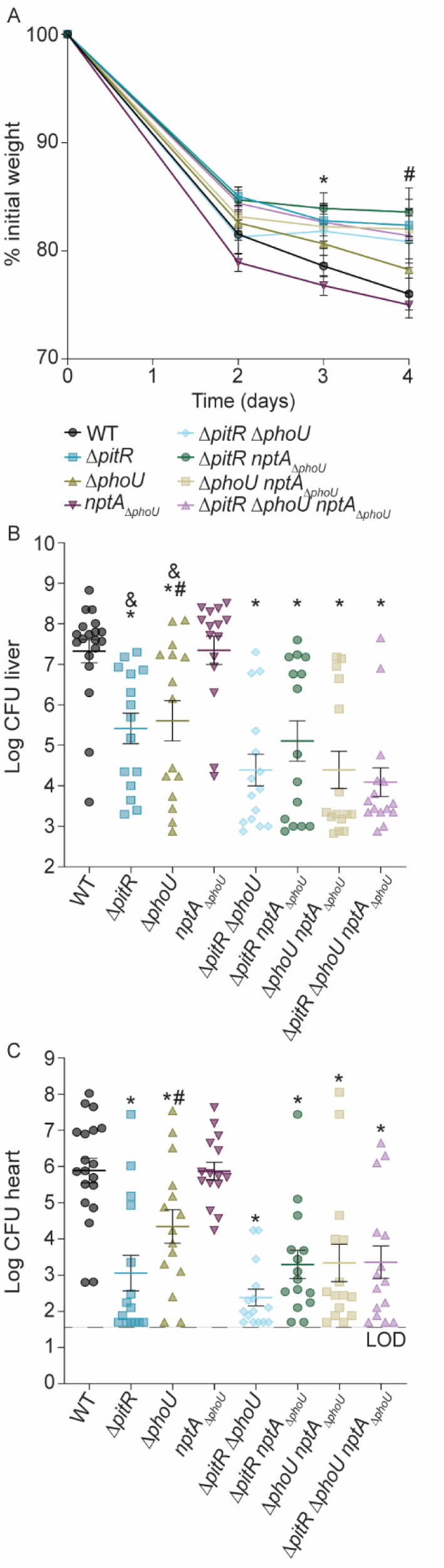
Multiple accessory proteins contribute to the ability of *Staphylococcus aureus* to cause infection. Wild-type C57BL/6J mice were systemically infected with *S. aureus* wild-type and the indicated strains and weight loss was monitored (A) and bacterial burdens in the liver (B) and heart (C) were enumerated 4 days post-infection by plating for colony forming units. (A) *\** = *P* ≤ 0.05 for Δ*pitRnptA*_Δ*phoU*_ mutant compared to the wild-type (Day 3 p.i.) and *#* = *P* ≤ 0.05 for Δ*pitR*, Δ*pitRnptA*_Δ*phoU*_, Δ*phoUnptA*_Δ*phoU*_, and Δ*pitR*Δ*phoUnptA*_Δ*phoU*_ mutants compared to the wild-type (Day 4 p.i.) via two-way ANOVA with Dunnett’s posttest. Error bars = SEM. (B & C) *\** = *P* < 0.05 relative to wild-type by Mann-Whitney test. *#* = *P* < 0.05 relative to Δ*phoU* by Mann-Whitney test. *&* = *P* < 0.05 relative to Δ*pitR*Δ*phoUnptA*_Δ*phoU*_ by Mann-Whitney test. Only significant *P* values are shown. Lines indicate medians. The data are results from three independent experiments. *n* ≥ 14 for each group.

**Figure 6.**
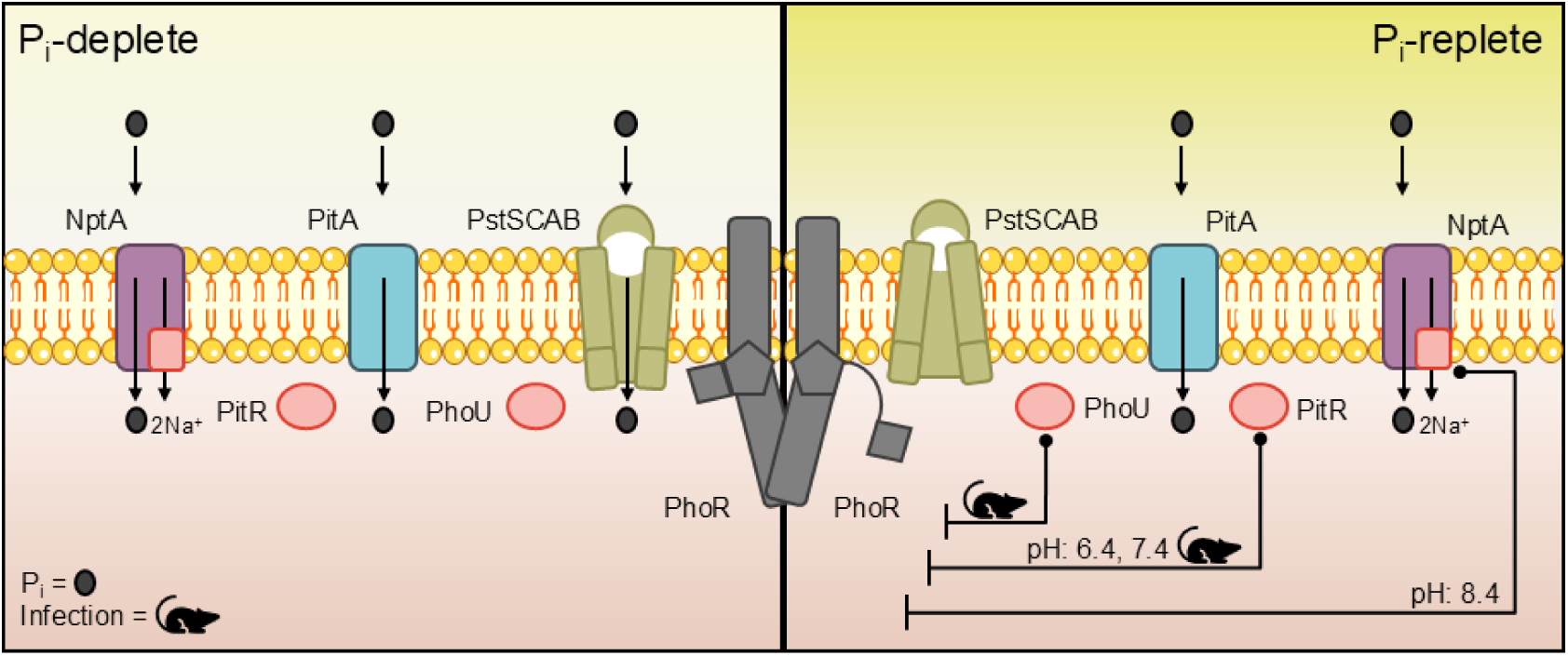
Model of the regulatory mechanisms controlling phosphate homeostasis. In P_i_-deplete environments, PhoR is in kinase conformation and P_i_ import is accomplished by the PstSCAB, PitA, and NptA transporters. In P_i_-replete environments, the accessory regulatory proteins exert repression of PhoR, which is in phosphatase conformation. The transporters PitA and NptA maintain P_i_ import activities, however PstSCAB expression and its subsequent P_i_ import is drastically decreased. The classical accessory regulatory protein PhoU associated with the PstSCAB P_i_-transporter represses PhoR *in vivo* during infection, along with the PitR protein associated with the PitA P_i_-transporter. In neutral and acidic conditions *in vitro*, PitR is the primary accessory regulatory protein that represses PhoR. In alkaline conditions *in vitro*, the PhoU-domain of NptA represses PhoR.

## Discussion

All organisms must tightly control phosphate uptake to balance the dual essentiality and toxicity of this nutrient (*1–18*). PhoU is critical to controlling P_i_ homeostasis and viability in *E. coli* and other microbes (*1, 4, 6–10, 28, 42, 43*). The current investigation revealed that bacteria possess at least three distinct groups of accessory regulators that control P_i_ homeostasis: the classical PhoU, PitR and the PhoU-domain of NptA, with a plurality of bacteria possessing two or more. Further, each clade regulates P_i_ homeostasis in conjunction with a specific family of P_i_-importers: PhoU with the Pst ABC-transporter family, PitR with the Pit family, and the NptA sodium-phosphate transporter family. The expanded repertoire of accessory regulators enables *S. aureus* to maintain P_i_ homeostasis in diverse environments with both PhoU and PitR independently contributing to infection. Cumulatively, these investigations reveal that bacteria possess a greater repertoire of P_i_ accessory regulators than previously appreciated, with each having critical non-redundant roles in regulating phosphate homeostasis and the ability of microbes to adapt to changing environments, including those encountered within the host.

Our mechanistic understanding of bacterial P_i_ homeostasis is primarily informed by *E. coli*, which is reliant on a single PhoU and PstSCAB, to repress PhoBR (*1, 25, 49*). However, the current work revealed that there is considerable diversity in both the number and type of accessory regulators and P_i_-transporters possessed by bacteria, with this diversity impacting P_i_ homeostasis. Significantly, in *S. aureus* PitR is the primary regulator of PhoPR activity under standard laboratory conditions. However, all three clades of accessory regulators contribute to the control of PhoPR and phosphate homeostasis. In *E. coli,* loss of PhoU results in a loss of viability (*4, 6*). It also led to the classical view that PhoU is primarily a regulator of the phosphate-responsive TCS. However, studies with *E. coli, S. pnenoniae* and *Sinorhizobium meliloti* suggest that PhoU also regulates the import activity of PstSCAB (*6, 9, 10*). In alkaline medium, where NptA transport is maximally active, the growth of a strain reliant on NptA_ΔPhoU,_ decreases as P_i_ concentration increases, raising the possibility that the alternative PhoU clades also modulate P_i_ import. Regardless of the mechanism, the current work suggests that PitRs and PhoU-domains of NptA contribute to controlling activation of the phosphate responsive TCS and P_i_ homeostasis.

In niches ranging from the stomach, skin and phagolysosome to the soil, changes in pH are a hurdle that all bacteria must be overcome (*44, 50–53*). While all three regulators can modulate the activity of PhoPR, the current work suggests a hierarchy that is influenced by environmental pH. In neutral and acidic media, PitR is the dominant regulator in *S. aureus*, with the PhoU-domain of NptA becoming more important in alkaline media. This corresponds to the environments in which PitA and NptA optimally import P_i_ (*31*). In *Clostridium acetobutyliculum*, which possesses PitR and PitA, PhoU and Pst systems (Data S1), alkalinization increases *pst* transcription (*54*). Most bacteria possess PitA and NptA homologs, with 533 (58%) of the sampled genomes possessing PitA and PitR, NptA with a PhoU-domain or both. With the current observations, this suggests that possessing multiple accessory regulators enables bacteria to maintain P_i_ homeostasis during environmental pH changes. However, it seems likely that pH is not the only environmental factor that modulates the contribution of the accessory regulators to controlling P_i_ homeostasis, as loss of PhoU by itself did not alter the activation of PhoPR in culture, but both Δ*pitR* and Δ*phoU* reduced the ability of *S. aureus* to cause infection. Similarly, the two *S. pneumoniae* PhoUs both independently contribute to infection (*55*).

Studies utilizing *E. coli* have led to a model in which phosphate sufficiency is sensed by PhoU monitoring translocation of P_i_ by PstSCAB (*1, 4, 6, 25–29*). However, the idea that bacteria are broadly reliant on the import of P_i_ as the sole means to evaluate P_i_ sufficiency and modulate P_i_ homeostasis is increasingly in doubt (*56–58*). In *S. enterica* Typhimurium, PhoBR activity is modulated by phosphate sources that bypass the Pst transporter or by altered magnesium levels (*56*). In *Ralstonia solanacearum*, the activity of the PhoB response regulator is modulated by PhoU in conjunction with PhoR or VsrB, the regulator of exopolysaccharides (*57*). In *B. subtilis,* which lacks any PhoU homolog (similar to 11% of the genomes examined), the activity of the response regulator of the phosphate-responsive TCS is modulated by lipoteichoic acid (LTA) intermediates (*59–62*). While in *Caulobacter crescentus,* which the current analysis indicates possesses only PhoU (Table S2), its loss does not alter PhoBR activity (*8*). Further detracting from the established model, in P_i_-replete medium *pst* expression in Δ*pitR*Δ*phoUnptA*_Δ*phoU*_ is not equivalent to that of wild-type bacteria or the triple mutant in P_i_-limited medium, suggesting the presence of additional modulators of PhoPR activity. However, if bacteria do not rely on import to sense P_i_ availability, it is unclear why there is a strong co-occurrence between P_i_-transporters and the associated accessory regulator. Further supporting the classical model of P_i_ sensing, in *S. aureus* loss of PitR leads to increased PhoPR activity without a change in total cellular P_i_. Similarly, growth in alkaline medium, which shifts P_i_ import away from PitA towards NptA (*31*), increases the regulatory contribution of the NptA PhoU-domain without altering intracellular P_i_ levels. Thus, the current work suggests, at least in *S. aureus*, that both P_i_ import and cellular sufficiency impact the activity of PhoPR. When combined with other work, it also suggests that the mechanism utilized by any given bacteria to regulate P_i_ homeostasis may lie on a spectrum somewhere between strictly relying on import and sensing cytoplasmic sufficiency signals. Given that P_i_ is an essential, but toxic nutrient, with a high cellular demand that is influenced by the environment and the physiological state of the cell, it is not surprising that bacteria have developed overlapping mechanisms to ensure sufficiency while limiting the potential for toxicity.

P_i_ homeostasis impacts the ability of bacteria to survive stressful situations and numerous links between P_i_ regulation and virulence have been identified (*1, 24, 39, 63*). Mutations that presumptively over activate the P_i_ starvation response reduced the virulence of many animal and plant pathogens (*64–70*), while in others, including *S. aureus,* preventing activation of this response reduces virulence (*35, 71–74*). As aberrant activation of the staphylococcal phosphate-responsive TCS reduces *S. aureus* virulence, this implies that pathogens encounter both P_i_ limitation and excess during infection. It also suggests that therapeutic perturbation of P_i_ homeostasis in either direction is likely to impair *S. aureus* virulence. Targeting the regulators of P_i_ homeostasis also has the potential to re-sensitize *S. aureus* and other microbes to currently approved therapeutics, as a mutation in the phosphate-responsive TCS and PhoU homologs are known to sensitize bacteria to multiple classes of antibiotics (*27, 33, 75–77*). The current work reveals that the expanded repertoire of accessory regulators and the associated complexity critically contributes to microbial success in diverse environments including within the host. Leveraging the potential vulnerability of microbial phosphate homeostasis to benefit human prosperity and health will require expanding the study of this process beyond model systems and developing a molecular understanding of how it functions.

## Materials and Methods

### Construction and analysis of the bacterial genome dataset

Annotations for 931 bacterial genomes were extracted from a hand-curated set, mostly derived from NMPDR and PATRIC (*78, 79*). Due to the history of this dataset, pathogenic bacteria are overrepresented. We excluded 3 genomes due to obvious poor quality, and 2 “synthetic genomes", leaving 926 genomes in this analysis. The bulk of the gene distribution analyses presented here are based upon proteins with the annotations listed in Table S1. Because sequencing and assembly errors are not easily distinguished from mutations, our analyses do not exclude proteins that might be defective. The occurrences of each annotation in each of the 926 genomes are compiled in Data S1.

### Sequence acquisition for PhoU-and PitR-related proteins

PhoU-related sequences were obtained by downloading all 36,026 entries from the PhoU Pfam domain class on InterPro. We limited the sequences to those with the same domain architecture as known PhoU proteins, e.g., in *E. coli* and *S. aureus*, which have two instances of the PhoU-domain (PF01895) with no additional known Pfam domains. Other PhoU-domain-containing proteins also encode other domains such as the Na_Pi_cotransporters (PF02690), which are not included in this analysis. Similarly, 20,380 sequences with the same domain architecture as the *S. aureus* PitR, which contains a single PhoU_div (PF01865) domain, were downloaded from InterPro.

### Gene tree analysis

To visualize the sequence relationship by a phylogeny, we further reduced the size of the protein sequences by first grouping each of the PhoU-and PitR-related protein sets by taxonomy at the genus level, selecting one representative sequence at random. Then, we randomly selected 200 and 100 sequences from the remaining ones for the two groups and added back the two *S. aureus* proteins. The resulting 302 protein sequences were aligned using MAFFT (v7) with the E-INS-i algorithm, which was optimized for distantly related proteins with multiple conserved domains. The aligned sequences were used as input for reconstructing a Maximum Likelihood (ML) tree using RAxML v8.2.12 with the LG+G substitution model. The best ML tree was shown as an unrooted phylogram with PhoU-and PitR-related sequences colored similarly as in the CLANS analysis.

### CLANS sequence clustering analysis

In CLANS, or CLuster Analysis of Sequences, biological sequences are compared using BLAST and represented as vertices in the graph, with edges representing high-scoring segment pairs (HSPs) from all-against-all BLAST. A variant of the Fruchterman and Reingold graph layout algorithm was run to generate a useful representation of pairwise sequence similarities, with attraction forces proportional to the negative logarithm of the HSP’s *P*-value. To perform the CLANS analysis, we first clustered the full PhoU-and PitR-related protein datasets by sequence similarity using CD-HIT (v4.8.1) at a threshold of 0.7 (70% identity). One sequence from each cluster was chosen at random as the representative, except for the two clusters containing the *S. aureus* proteins, where *S. aureus* sequences were manually selected (SaPhoU: A0A0H3K808, SaPitR: A0A0H3K7D8). This reduced the number of sequences to 7,933 and 4,672 for the two groups, respectively. The filtered set of sequences were combined and submitted to the CLANS web app on the MPI Bioinformatics Toolkit server (https://toolkit.tuebingen.mpg.de/tools/clans), which performed the all-against-all BLASTp and returned a .clans format result. To visualize the result, we used the clans.jar Java app to open the result, set the cooling parameter to 0.8 and used default parameters for the rest, important ones include {attraction: 10; attraction exponent: 1; repulsive: 10, repulse exponent: 1}. The algorithm was run till convergence. PhoU-and PitR-related proteins were grouped and shown by different colors. A 2D-cluster was shown for visualization.

### Bacterial strains

For routine culture, *Staphylococcus aureus* was grown in tryptic soy broth (TSB) and on tryptic soy agar (TSA) plates, while *Escherichia coli* strains were cultivated in Luria broth (LB) and on Luria agar plates. Both species were grown at 37°C. As needed, 100 μg/mL of ampicillin or 10 μg/mL of trimethoprim for *E. coli* and 10 μg/mL of chloramphenicol or 10 μg/mL of trimethoprim for *S. aureus* was added to the medium. Bacterial stocks were stored at −80°C in brain-heart infusion (BHI) broth supplemented with 30% glycerol. A complete list of strains and plasmids are listed in Tables S2 & S3.

### Mutant construction

For the generation of the *S. aureus* Δ*pitR* and Δ*phoU* mutants, the 5′-and 3′-flanking regions (∼1 kb upstream and downstream) of the target gene were amplified using the indicated primers (Table S4). These fragments were cloned into pKOR1 via site-specific recombination using the Gateway BP Clonase II enzyme mix (Thermo Fisher Scientific) (*35*). *S. aureus* Δ*pitR*Δ*phoU* was generated by transducing the *phoU::erm* allele into Δ*pitR* via Phi85. To generate the *nptA*_Δ*phoU*_, Δ*pitRnptA*_Δ*phoU*_, Δ*phoUnptA*_Δ*phoU*_, Δ*pitR*Δ*phoUnptA*_Δ*phoU*_ single, double and triple mutants pKOR1::*nptA*_Δ*phoU*_ was used in conjunction with wild-type, Δ*pitR,* Δ*phoU,* and Δ*pitR*Δ*phoU*, respectively. This plasmid was created by amplifying approximately 1 kb of DNA upstream and downstream of the encoded PhoU-domain of *nptA* and inserting the fragments into pKOR1 using the Gibson Assembly Master Mix (New England BioLabs), thereby removing the entire PhoU-domain (residues 450-534) of *nptA* while retaining the stop codon. Complementation constructs for *pitR* and *nptA* were created by cloning the coding sequence and 5’ intergenic regions into pKK30, a low copy plasmid, using the indicated primers (Table S4). The *phoU* gene was also cloned into pKK30 but under the control of the constitutive p*lgt* promoter (*35*). Plasmids were electroporated into *S. aureus* RN4220 and then transferred into the final recipient strain. All constructs were verified by sequencing and each staphylococcal mutant was confirmed to be hemolytic on blood agar plates.

### Growth assays and expression analysis

Growth and expression assays utilized a defined medium, PFM9, with assays performed as previously described (*31, 35*). Briefly, phosphate-free M9 salts were supplemented with amino acids, vitamins, 0.5% glucose, 2 mM MgSO_4_, 1 mM CaCl_2_, 1 µM FeSO_4_, ZnSO_4_, and MnCl_2_. The pH of the medium was manipulated by the addition of 70 mM MOPS (morpholinepropanesulfonic acid) for pH 6.4, HEPES (N-2-hydroxyethylpiperazine-N-2-ethane sulfonic acid) for pH 7.4, and Tris for pH 8.4. A 10X P_i_ source stock composed of KH_2_PO_4_ and Na_2_HPO_4_ was added to the medium to achieve the desired P_i_ concentration. For these assays, a bacterial colony was inoculated into 5 mL TSB in a 15 mL conical tube and grown for 8 h at 37°C on a rollerdrum. Using a 15 mL conical tube, the culture was then back-diluted 1:10 into 5 mL PFM9 plus 70 mM HEPES (pH 7.4) containing 158 µM P_i_ and grown for 16h at 37°C on a roller drum. Overnight cultures were inoculated at a 1:100 dilution into a 96-well round-bottom plate containing 100 µl/well PFM9 and incubated at 37°C with shaking at 180 rpm. The optical density at 600 nm (OD_600_) was measured to monitor bacterial growth. Expression or RFU (relative fluorescence units) was determined by measuring fluorescence (excitation and emission wavelengths of 505/535 nm), normalizing the values to the OD_600_.

### Phosphate accumulation assays

Phosphate accumulation assays were performed largely as previously described (*31*). Bacteria were pre-cultured in PFM9 as described above for phosphate growth assays, with the exception that the bacteria were cultured in 50 mL conical tubes containing 15 mL of medium. The bacteria were grown to an OD_600_ of ∼0.25-0.3 before harvesting. The cells were washed once with 5 mL Tris-EDTA (TE) buffer and then lysed in 500 µl TE buffer by mechanical disruption. Particulate matter was removed by centrifugation. The total protein concentration measured by the Pierce bicinchoninic acid (BCA) assay kit was utilized to normalize the P_i_ concentration. The P_i_ concentration was measured with the Biomol Green kit (Enzo Life Sciences) according to the manufacturer’s instructions following the treatment of the cell lysate with yeast exopolyphosphatase.

### Animal infections

Mouse infections were performed as previously described, with minor modifications (*31, 35, 80*). Briefly, a bacterial colony of wild-type and PhoU homolog mutant strains of *S. aureus* were inoculated into 5 mL TSB in a 15 mL conical tube and grown for 16 h at 37°C. The culture was then back-diluted 1:100 into 5 mL TSB and grown for 3 h at 37°C on a roller drum. The bacteria were washed and resuspended in phosphate-free, bicarbonate-buffered saline (BBS) and diluted to a density of 10^8^ CFU/mL. Ten-week-old female C57BL/6J mice were retro-orbitally injected with 10^7^ CFU in 100 µl of buffer. The infection was allowed to proceed for 96 h before the mice were sacrificed. The livers, hearts, and kidneys were harvested. Organs were homogenized and bacterial burdens were determined by plating serial dilutions.

## Author contributions

Conceptualization: CV, TKF

Data curation: CV, GJO, BZH, TKF

Formal analysis: CV, GJO, BZH, TKF

Funding acquisition: TKF, BZH

Investigation: CV, GJO, BZH, ESJS, JNR

Methodology: CV, GJO, BZH

Visualization: CV, BH, TKF

Supervision: TKF

Writing—original draft: CV

Writing—review & editing: CV, GJO, BZH, TKF

## sData and materials availability

All data needed to evaluate the conclusions of this paper are present in the paper and/or the Supplemental Materials. The MPI Bioinformatics Toolkit server is available at https://toolkit.tuebingen.mpg.de/tools/clans. Bacterial genomes from the paper can be derived from NMPDR and PATRIC at https://www.bv-brc.org/. Sequences from the paper can be obtained from InterPro at https://www.ebi.ac.uk/interpro/.

